# Soil water consumption characteristics in rain-fed apple orchards and wheat fields

**DOI:** 10.1101/2022.04.08.487632

**Authors:** Lu Zhang, Yiquan Wang, Zenghui Sun

## Abstract

Agricultural production in Weibei rain-fed highland, Northwest China, is facing severe drought and water shortages. Here, soil water consumption characteristics in rain-fed orchards and farmlands were explored to ascertain the rationality of planting orchards in Weibei. Soil moisture dynamics was monitored in the 0–150-cm soil profiles of different aged ‘Red Fuji’ apple orchards (young: 7 years, mature: 13 years, and old: 22 years), and in long-term cultivated winter wheat fields during the growing season of apple trees. The over-consumption and consumption of soil water were analyzed to evaluate water stress and differential water consumption by distinct vegetation, respectively. Soil desiccation index was used to determine the occurrence of dry soil layers. Generally, there was no water stress in the 0–150-cm orchard soil profiles, while water stress was observed at the 0–70-cm soil depths in the old orchards (mid-June) and farmlands (mid-May–mid-July). Water consumption took place at deeper depths for longer periods in the older orchards than in the younger orchards. Soil desiccation was not observed in the young orchards, while mild desiccation occurred at the 0–80-cm soil depths in the mature and old orchards in mid-June. The desiccation intensity was mild at the 0–60-cm soil depths in mid-April–mid-May, intense at the soil 0–150-cm depths in mid-June, and moderate at the 20–150-cm soil depths in mid-July. In conclusion, conversion from wheat fields to apple orchards could reduce soil water stress, reduce dry soil layers, and mitigate soil desiccation in the rain-fed highland area.

## Introduction

Soil water regime influences regional land use, vegetation layout planning, and ecological construction. Generally, rain-fed highlands are characterized by inadequate moisture conditions and poor irrigation facilities [1-4], while having abundant light and heat resources [5-7]. Weibei rain-fed highland is a critical agricultural region in Northwest China, where water shortage is the primary factor limiting sustainable crop production. Since China implemented the reform and opening-up policy and adjusted agricultural production structure, the land use pattern in Weibei has shifted gradually from crop production to orchard farming [8]. Such conversion of land use aims to address the shortcomings of regional resources and take advantage of the regional natural resources, to transform them into economic resources and improve the people’s livelihoods. However, our understanding of the influence of fruit industry development on regional water resources and the rationality of planting orchards in Weibei over the past four decades remains poor.

Many studies have evaluated soil moisture content based on the physiological responses of crops [9-13]. Ковалев (1981) argued that vegetation not only consumes but also conserves soil water [14]. In the absence of significant changes in natural conditions (e.g., climate and soil), land cover largely influences the occurrence and severity of soil drought [15-18]. Studies have suggested that soil water storage capacity in vegetated land is substantially higher than in bare or fallow land [19-21]. In addition, the spatiotemporal distribution and migration of soil water under different land cover types have been extensively studied [22-28]. Yang et al. [29] reported that different intensities of desiccation occurred below 3-m soil depths in various forestlands, excluding young apple orchards and shrub stands (*Hippophae rhamnoides* and *Caragana korshinskii*); soil desiccation intensity increased with an increase in the age of forest trees. Tian et al. [30] also observed that afforestation enhanced water supply in the 0–3-m soil profile of forestlands in loess hilly areas, despite the occurrence of dry soil layers. Bai et al. [31] observed distinct spatial and seasonal soil water trends in the Qilian Mountains, which were closely related to different land cover types. To date, few comparative studies have explored soil moisture dynamics and water consumption characteristics in Weibei region following the conversion of the major vegetation from annual winter wheat to perennial fruit trees.

According to Ковалев (1981), when evaporation is dominant (equivalent to the rate in bare land after crop harvesting), soil water consumption occurs mainly in the upper soil layers; however, when transpiration is dominant (equivalent to the rate in vegetated land during crop growing seasons), soil water consumption is concentrated mainly in the soil layers with dense roots [14]. The findings suggest that the location and intensity of soil desiccation vary under different water consumption patterns across growing seasons of various vegetation types. Here, we monitored the spatiotemporal dynamics of soil moisture content and evaluated soil water consumption in Weibei following conversion of farmland into orchards, to examine the rationality of planting orchards in the region considering the available water resources.

## Materials and Methods

### Sites Description

The study was conducted in Zhanghong Town, a typical rain-fed highland area in Weibei (E 108°08′–108°52′, N 34°57′–35°33′), located in the central part of Xunyi County, Shaanxi Province, China (**Figure 1**). The study area falls under the fruit eugenic development area in the Weibei loessial rain-fed highland. The mean elevation is the highest in Weibei, 1155 m a.s.l. The terrain is complex, with higher relief in the northeast and lower relief in the southwest. The landform exhibits typical hilly and gully features of the Loess Plateau. The mean annual temperature in the region is 9.1°C. The coldest month is January, with the lowest temperature being -24.3°C; while the hottest month is July, with the highest temperature being 36.3°C. The annual temperature difference is –26°C, and the mean frost-free period is approximately 180 days.

**Fig 1.**
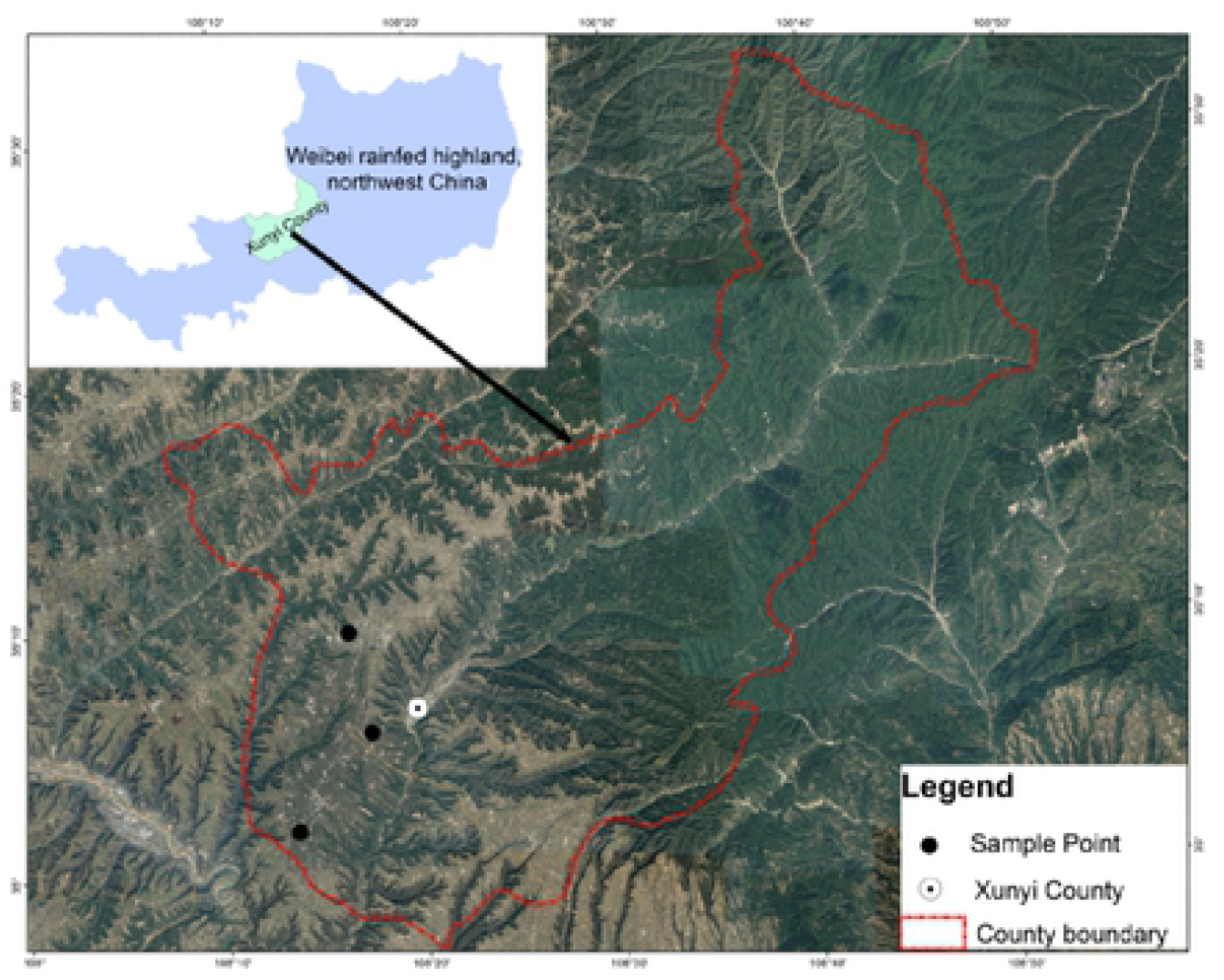
Map of the study area in Xunyi County, Shaanxi Province, Northwest China.

Xunyi has abundant sunlight resources, with an annual total solar radiation of up to >500 kJ/cm^2^ and annual total sunshine hours >2300 h; the sunlight resources are most abundant in May and June. The considerable diurnal and annual temperature differences are favorable for sugar accumulation in apples and high-quality fruit production. In the year of the study (2019), the mean monthly temperature in Xunyi was 6.4°C in March (sprouting in spring) and 15.4°C in September (harvesting in fall). The >0°C cumulative temperature and ≥10°C effective cumulative temperatures in the year were 3834.5°C and 3534.1°C, respectively.

### Soil Sampling

Apple (*Malus domestica* Borkh. cv. Red Fuji) orchards of different ages (young: 7 years, mature: 13 years, and old: 22 years) were selected in Zhanghong Town, with three orchards per group. Apple trees were planted 2 m apart with rows 4 m apart in each orchard. Winter wheat (*Triticum aestivum* L. cv. Xiaoyan 22) fields 20 cm apart were selected in the vicinity of orchards as controls, with one plot per group. The plot sizes of orchards and farmlands were 20,000 m^2^ (500 m [length] × 400 m [width]).

Soil moisture content was monitored in each plot in an apple tree-growing season, namely, from the beginning of March, when apple trees sprouted, to the end of September, when apple fruits were harvested. Soil sampling was conducted monthly (seven times in total). During each sampling activity, a 6-cm-diameter soil augur was used to collect soil samples at eight depth intervals in the 0–150-cm profile (0–10, 10–20, 20–40, 40–60, 60–80, 80–100, 100–120, and 120–150). In the orchard plots, four sampling points were selected 1 m away from the trunks of five random trees in four directions symmetrically, and the four samples from each tree were mixed to form a composite sample. In the farmland plots, soil samples were obtained at five points based on the diagonal method (one at the central point of the plot and four at the mid-point of the straight lines connecting the central point to the four corners).

### Data Analysis

Soil water consumption and relevant parameters were calculated based on depth and month using Equations. (1)–(3).

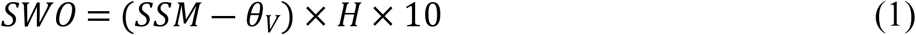

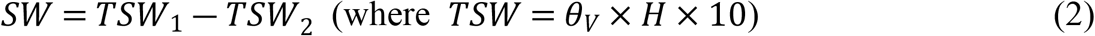

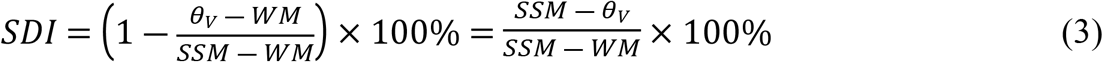

In Eq. (1): *SWO* is soil water over-consumption, mm; *θ*_v_ is soil moisture content by volume, %; *SSM* is stable soil moisture content (arithmetic mean of field capacity and wilting moisture content), %; and *H* is soil depth divided by 10 (*H* = 1, 2, 3…).

In Eq. (2): *SW* is soil water consumption, mm; *TSW*_1_ is soil water storage at the precedent sampling time *t*_1_, mm; and *TSW*_2_ is soil water storage at the next sampling time *t*_2_, mm

In Eq. (3): *SDI* is soil desiccation index, %.

Based on the *SDI* value, soil desiccation intensity could be divided into the following six levels: ① *SDI* < 0, no desiccation; ② 0 ≤ *SDI* < 25%, mild desiccation; ③ 25% ≤ *SDI* < 50%, moderate desiccation; ④ 50% ≤ *SDI* < 75%, severe desiccation; ⑤ 75% ≤ *SDI* < 100%, intense desiccation; and ⑥ *SDI* ≥ 100%, extreme desiccation. Levels ② –⑥ indicate the occurrence of mildly, moderately, severely, intensely, and extremely dry soil layers, respectively [32].

Data are means ± standard deviation (*n* = 5). Statistical analyses were performed using the t-test in IBM SPSS Statistics 20.0 (IBM Corp., Armonk, NY, USA). A *P* value less than 0.05 was considered to indicate statistical significance.

## Results

### Profile Distribution of Soil Moisture

The distribution characteristics of soil moisture in the 0–150-cm profile of orchards and farmlands are illustrated in **Figure 2**. With increasing soil depth, soil moisture content was variable and substantially higher in orchards than in local farmlands. The mean soil moisture contents in the profiles of young, mature, and old orchards were 17.6%, 18.5%, and 14.1% higher than the contents in the farmlands, respectively. The minimum soil moisture contents in the profiles of young, mature, and old orchards were 8.1%, 8.7% and 6.4% higher than local stable soil moisture content, respectively. Conversely, the soil moisture content at the 0–60-cm depths of farmland was 5.81% lower than the local stable moisture content. However, soil moisture content exhibited increasing trends in farmlands with increases in soil depth; it exceeded the local stable moisture content at 60-cm depth and then leveled off in deeper soil layers. The mean soil moisture contents in the profiles of different orchards were in the following order: mature orchards > young orchards > old orchards. Specifically, the mean soil moisture contents in the mature orchards were 0.8% and 3.9% higher than the contents in the young and old orchards, respectively, and the mean contents in the young orchards were 3.1% higher than the contents in the old orchards.

**Fig 2.**
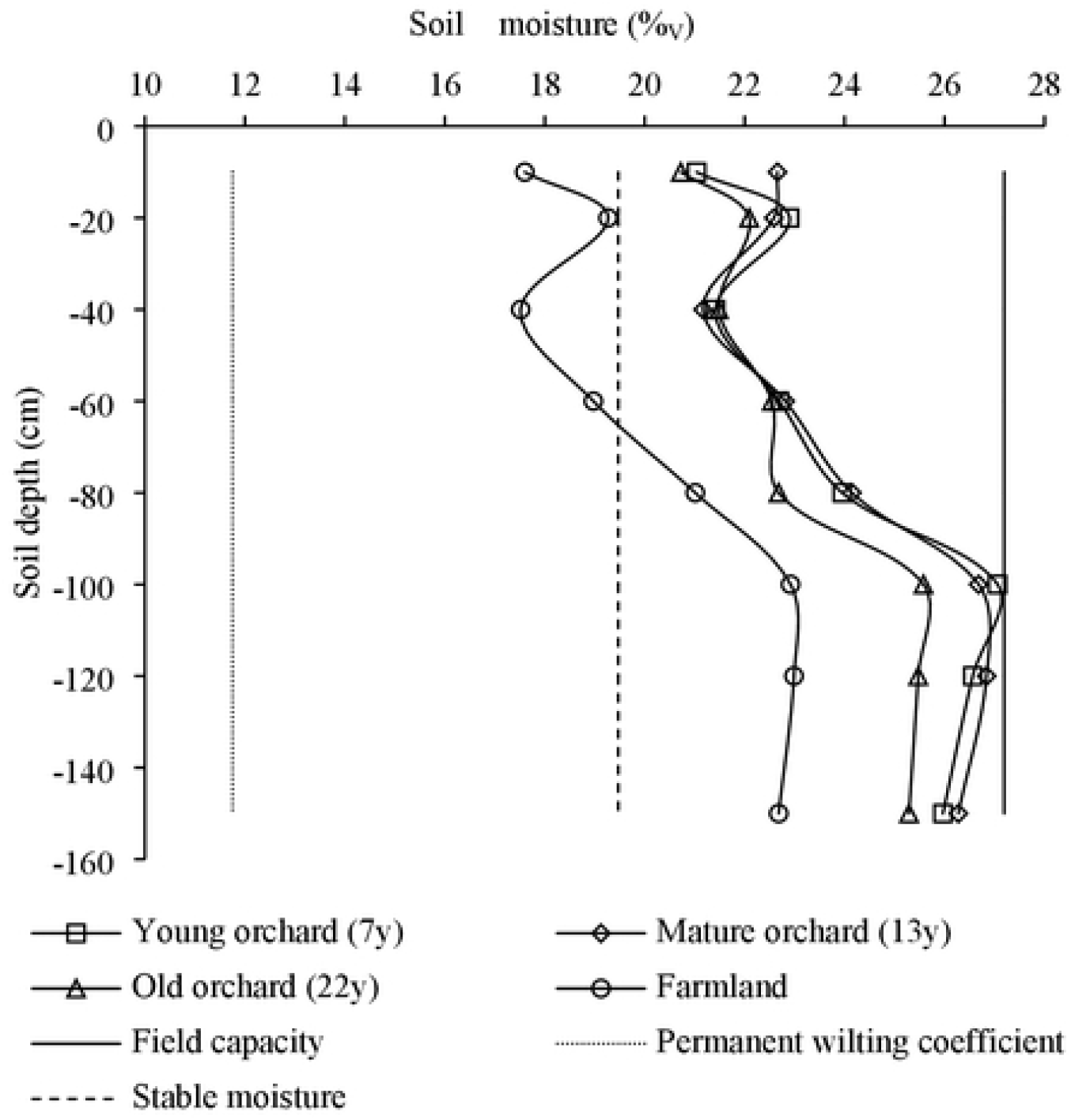
Soil moisture distribution in the 0–150-cm profile in apple orchards and farmlands.

### Differences in Soil Water Over-consumption

The soil water over-consumption estimates were negative values with minor differences between groups in most cases, and positive values were obtained from the old orchards only in mid-June (**Figure 3**). During the apple-growing season, the mean soil moisture contents in the 0–150-cm profiles of young, mature, and old orchards were 23.0%, 24.0%, and 19.3% higher than the local stable moisture content, respectively. In addition, positive soil water over-consumption values were obtained from farmlands in mid-May, mid-June, and mid-July; the highest over-consumption occurred in mid-June, which was 9.4-fold that of mid-May and 1.6-fold that of mid-July. The mean soil moisture content in the farmland soil profiles was 4.9% higher than the local stable moisture content. In addition, there were significant differences in soil water over-consumption between the orchards and farmlands across different periods (*P* <0.05).

**Fig 3.**
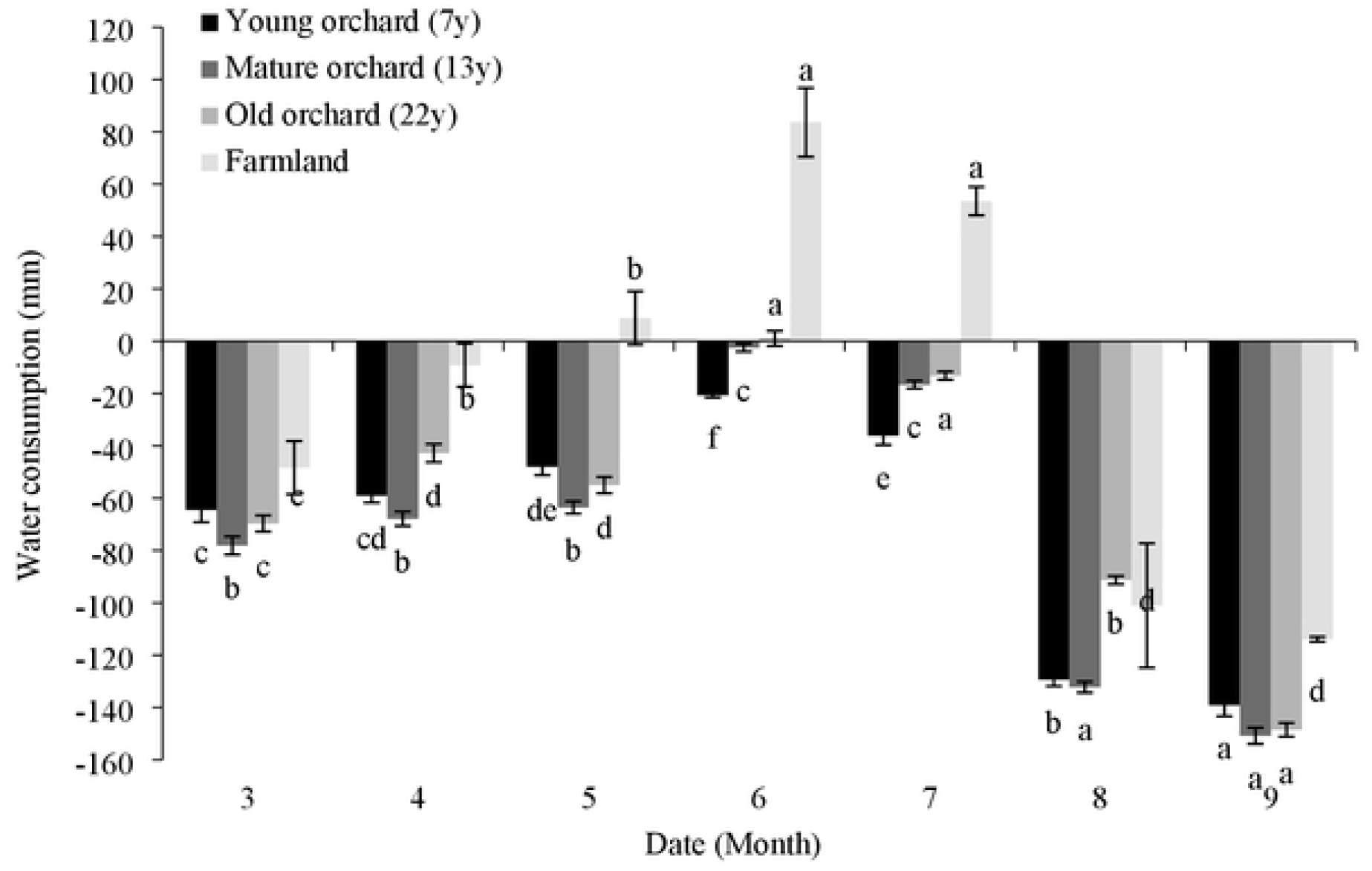
Soil moisture consumption in apple orchards and farmlands in the course of an apple-growing season. Lower case letters above or below the error bars indicate significant differences between groups (*P* < 0.05).

### Soil Water Surplus

Soil water consumption in the 0–150-cm soil profile occurred in the young and mature orchards in mid-May to mid-June, while in the old orchards and farmlands, water consumption occurred mainly in mid-March to mid-April, and mid-May to mid-June (**Table 1**). Soil water consumption took place at deeper depths for longer periods in farmlands when compared to orchards. The total water consumption in the soil profile was ranked as follows: old orchards > farmlands > mature orchards > young orchards. Soil water surplus in the young orchards relative to the mature and old orchards was 2.7% and 467.9%, respectively, while the soil water surplus in the mature orchards and farmlands relative to the old orchards was 453.1% and 398.8%, respectively.

**Table 1.** Water consumption in apple orchard and farmland soil profiles (mm). † Lower case letters in the same column indicate significant differences between different groups (*P* < 0.05).

| Crop | Period | Soil depth (cm) |  |  |  |  |  |  |  |  |
| --- | --- | --- | --- | --- | --- | --- | --- | --- | --- | --- |
|  |  | 0–10 | 10–20 | 20–40 | 40–60 | 60–80 | 80–100 | 100–120 | 120–150 | Σ |
| Young orchards | 3.15–4.15 | 3.16±1.33<br>a<sup>†</sup> | 0.89±0.37<br>b | 2.55±0.58<br>b | 1.59±0.95a | -0.61±0.7<br>9b | -1.12±0.21<br>a | -0.60±0.2<br>5a | -0.50±2.1<br>0a | 5.36±0.12a |
|  | 4.16–5.15 | -6.10±0.6<br>2a | -1.15±0.4<br>4a | -1.42±1.4<br>6a | 2.43±1.17<br>b | 5.74±1.69<br>a | 5.41±1.22a | 4.54±0.91<br>a | 1.63±0.75<br>a | 11.08±1.76<br>a |
|  | 5.16–6.15 | 3.33±1.28<br>a | 2.18±0.08<br>b | 5.91±0.24<br>a | 6.60±0.29a | 1.86±0.23<br>a | 0.28±2.42a | 0.48±2.48<br>b | 6.75±2.21<br>a | 27.39±7.77<br>a |
|  | 6.16–7.15 | -7.68±0.1<br>6a | -2.68±0.8<br>0a | 0.78±1.36<br>a | -1.59±2.74<br>a | -1.40±1.1<br>3a | -0.50±1.40<br>a | 0.21±1.84<br>a | -2.61±2.0<br>1a | -15.46±0.76<br>a |
|  | 7.16–8.15 | -4.80±1.0<br>2a | -6.81±0.7<br>4ab | -18.05±1.<br>91a | -18.45±4.1<br>5a | -15.89±0.<br>18a | -13.39±0.7<br>3ab | -12.67±2.<br>78a | -3.18±2.2<br>9a | -93.24±2.19<br>a |
|  | 8.16–9.15 | 1.70±0.13<br>ab | 1.72±0.36<br>ab | 2.31±0.27<br>b | 1.97±0.67a | 1.86±0.52<br>ab | -1.04±0.36<br>a | -1.19±3.1<br>0a | -17.09±2.<br>13a | -9.76±4.39a |
|  | Σ | -10.38±0.<br>39b | -5.85±1.0<br>8b | -7.92±0.3<br>7ab | -7.45±0.80<br>a | -8.44±0.4<br>8a | -10.36±0.2<br>3a | -9.22±0.3<br>7a | -15.00±1.<br>73a | -74.62±0.15<br>a |
|  | % | 21.7 |  | 10.6 | 10.0 | 11.3 | 13.9 | 12.4 | 20.1 | 100 |
| Mature orchards | 3.15–4.15 | 3.25±0.68<br>b | 3.04±0.42<br>c | 2.80±0.65<br>c | 4.29±0.51a | 1.16±2.69<br>b | -2.56±2.03<br>a | -1.37±1.2<br>7a | -0.35±2.1<br>7a | 10.25±7.04<br>a |
|  | 4.16–5.15 | -3.41±0.6<br>7a | -2.98±1.0<br>1a | 1.58±1.60<br>a | 3.53±1.10<br>b | 2.45±0.97<br>a | 4.05±1.51a | 3.14±1.00<br>a | -4.05±0.3<br>5b | 4.31±0.88a<br>b |
|  | 5.16–6.15 | 6.37±1.94<br>a | 6.95±2.11<br>a | 12.78±3.7<br>8a | 5.82±3.53a | 5.34±4.46<br>a | 6.50±4.16a | 5.53±3.04<br>ab | 11.76±5.0<br>1a | 61.05±20.8<br>2a |
|  | 6.16–7.15 | -7.36±1.6<br>6a | -6.64±1.5<br>4ab | -4.58±3.5<br>8a | 2.35±4.10a | 1.68±4.28<br>a | 1.68±3.14a | -1.77±3.2<br>1a | 0.54±2.19<br>a | -14.10±20.4<br>4a |
|  | 7.16–8.15 | -5.77±1.1<br>8a | -4.97±0.8<br>7a | -21.37±2.<br>79a | -27.43±1.4<br>6ab | -21.84±0.<br>87a | -17.27±1.1<br>4ab | -9.17±2.5<br>8a | -7.77±5.4<br>1a | -115.60±0.7<br>0ab |
|  | 8.16–9.15 | 2.16±0.76<br>ab | 1.24±1.16<br>ab | 2.10±0.10<br>b | 3.87±0.58a | 3.69±0.68<br>a | -2.79±0.04<br>a | -8.98±3.3<br>0a | -19.86±4.<br>39a | -18.59±9.39<br>a |
|  | Σ | -4.77±1.2<br>8a | -3.38±0.4<br>6a | -6.69±1.5<br>9a | -7.56±1.27<br>a | -7.53±2.4<br>2a | -10.39±2.0<br>2a | -12.63±0.<br>56a | -19.73±0.<br>18a | -72.68±0.59<br>a |
|  | % | 11.2 |  | 9.2 | 10.4 | 10.4 | 14.3 | 17.4 | 27.2 | 100 |
| Old orchards | 3.15–4.15 | 3.28±1.41<br>a | 4.65±1.15<br>a | 2.87±2.05<br>b | 3.98±1.27a | 5.49±0.70<br>a | 1.40±0.15a | 1.88±0.18<br>a | 3.35±1.60<br>a | 26.88±3.30<br>a |
|  | 4.16–5.15 | -3.49±3.2<br>0a | -3.26±2.1<br>6a | -3.69±1.0<br>4a | -0.13±1.00<br>b | 1.86±0.73<br>a | 1.38±1.18a | -2.48±1.8<br>9b | -2.41±1.2<br>6b | -12.22±1.75<br>b |
|  | 5.16–6.15 | 2.08±3.75<br>a | 3.22±1.48<br>ab | 10.38±2.1<br>9a | 6.70±0.75a | 4.60±0.72<br>a | 8.88±2.64a | 9.44±3.27<br>ab | 10.77±3.9<br>7a | 56.06±8.33<br>a |
|  | 6.16–7.15 | -6.96±2.9<br>1a | -4.03±0.8<br>5ab | -2.86±1.9<br>8a | 3.59±0.46a | -0.39±1.5<br>2a | -2.06±3.54<br>a | -1.22±3.8<br>4a | -0.31±5.5<br>4a | -14.24±12.2<br>0a |
|  | 7.16–8.15 | -5.22±0.3<br>8a | -6.04±0.5<br>7ab | -20.55±3.<br>07a | -24.42±2.4<br>2ab | -14.31±4.<br>42a | -6.05±5.55<br>a | -1.20±2.8<br>9a | -0.28±3.3<br>6a | -78.09±21.9<br>0a |
|  | 8.16–9.15 | 0.79±0.85<br>b | 0.47±0.22<br>b | 2.26±0.67<br>b | 2.51±0.98a | -3.44±2.6<br>3b | -13.37±2.2<br>0b | -17.25±1.<br>57a | -29.19±2.<br>05a | -57.21±9.96<br>a |
|  | Σ | -1.59±0.2<br>5b | -0.83±0.7<br>2ab | -1.93±0.7<br>5b | -1.30±0.38<br>a | -1.03±0.9<br>9a | -1.64±1.41<br>a | -1.81±0.9<br>8a | -3.01±0.0<br>9a | -13.14±3.05<br>a |
|  | % | 18.4 |  | 14.7 | 9.9 | 7.9 | 12.5 | 13.7 | 22.9 | 100 |
| Farmlands | 3.15–4.15 | 6.95±1.34<br>a | 6.00±0.73<br>a | 10.08±0.9<br>2a | 4.12±1.02a | 0.25±1.52<br>b | 1.80±2.78a | 4.34±5.15<br>a | 5.66±7.66<br>a | 39.21±18.4<br>4a |
|  | 4.16–5.15 | -7.36±1.2<br>6a | -2.58±0.5<br>8a | 3.27±3.57<br>a | 4.52±1.86a | 6.52±2.91<br>a | 5.75±1.59a | 3.28±1.78<br>a | 4.61±1.30<br>a | 18.01±12.4<br>4a |
|  | 5.16–6.15 | 5.99±0.48<br>a | 6.05±0.73<br>ab | 8.15±2.35<br>a | 8.64±3.11a | 9.40±2.89<br>a | 11.34±3.4<br>4a | 11.65±3.8<br>4a | 13.54±5.4<br>7a | 74.77±21.2<br>1a |
|  | 6.16–7.15 | -7.73±1.5<br>9a | -7.85±1.7<br>1b | -7.30±2.5<br>4a | 0.05±2.31a | -0.07±1.0<br>4a | -1.34±1.80<br>a | -2.94±2.0<br>0a | -2.96±3.1<br>4a | -30.13±14.5<br>1a |
|  | 7.16–8.15 | -6.94±0.8<br>8a | -9.34±1.5<br>3b | -26.49±2.<br>95a | -32.29±2.3<br>0b | -30.41±1.<br>00b | -24.49±4.4<br>3b | -14.30±7.<br>89a | -10.25±9.<br>69a | -154.52±21.<br>82b |
|  | 8.16–9.15 | 4.24±1.20<br>a | 2.74±0.34<br>a | 6.48±0.97<br>a | 3.72±0.98a | 4.40±1.91<br>a | -1.69±5.38<br>a | -9.45±9.1<br>0a | -23.32±9.<br>16a | -12.88±23.5<br>4a |
|  | Σ | -4.84±0.4<br>6a | -4.98±0.2<br>8ab | -5.80±1.5<br>1a | -11.24±2.3<br>7a | -9.90±0.2<br>4a | -8.63±1.94<br>a | -7.44±3.3<br>8a | -12.71±8.<br>51a | -65.54±10.6<br>5a |
|  | % | 15.0 |  | 8.9 | 17.2 | 15.1 | 13.2 | 11.4 | 19.4 | 100 |

From mid-March, there was a surplus of soil water at the 60–80-cm depth in the young orchards, while soil water increased relatively at the 80–100-cm depth in the mature orchards. In the old orchards and farmlands, a surplus occurred at the 0–10 cm depth from mid-April. Considering the entire 0–150-cm profile, the highest total water consumption was observed from mid-May through mid-June in both orchards and farmlands. The total water consumption in the mature orchards increased by 122.9% and 8.9% when compared to the consumption in the young and old orchards, respectively, was although it reduced by 18.3% when compared to the level in farmlands. The lowest total water consumption and the highest water surplus occurred in mid-July to mid-August in orchards and farmlands.

The vertical distribution of soil water consumption varied based on orchard age. The highest water consumption rates were observed at the 40–60-cm depth in the young orchards, the 20–40-cm depth in the mature orchards, and farmlands, and at the 60–80-cm depth in the old orchards. The lowest water consumption rates were observed at the 0–20-cm depth in the young orchards, and the 120–150-cm depth in the mature orchards, old orchards, and farmlands.

### Soil Desiccation Intensity and Dry Soil Thickness

**Table 2** lists the spatiotemporal changes in soil desiccation intensity in orchards and farmlands. Considering the mean moisture content in the soil profile at a particular time, soil desiccation did not occur in the young orchards throughout the growing season of apple trees. However, mild soil desiccation was observed in the mature and old orchards in mid-June only.

**Table 2.** Soil desiccation intensity and dry soil thickness in orchards and farmlands in Weibei.

| Crop | Time (month) | SDI (%) | Desiccation intensity | Soil depth (cm) |  |  |  |  | Total thickness of dry soil layer (cm) |
| --- | --- | --- | --- | --- | --- | --- | --- | --- | --- |
|  |  |  |  | Extreme | Intense | Severe | Moderate | Mild |  |
| Young orchards | 3 | -46.8 | \ | \ | \ | \ | \ | 0–10 | 10 |
|  | 4 | -38.5 | \ | \ | \ | 0–10 | \ | \ | 10 |
|  | 5 | -37.1 | \ | \ | \ | \ | \ | \ | 0 |
|  | 6 | -11.7 | \ | \ | \ | \ | 0–10, 40–60 | 20–40, 60–80 | 70 |
|  | 7 | -32.8 | \ | \ | \ | \ | 40–60 | 20–40 | 40 |
|  | 8 | -117.3 | \ | \ | \ | \ | \ | \ | 0 |
|  | 9 | -117.0 | \ | \ | \ | \ | \ | \ | 0 |
| Mature orchards | 3 | -64.4 | \ | \ | \ | \ | \ | \ | 0 |
|  | 4 | -50.1 | \ | \ | \ | \ | \ | \ | 0 |
|  | 5 | -51.4 | \ | \ | \ | \ | \ | 40–60 | 20 |
|  | 6 | 6.5 | Mild | \ | \ | 20–40 | 0–20, 40–60 | 60–80 | 80 |
|  | 7 | -17.9 | \ | \ | \ | 40–60 | 20–40 | 60–80 | 60 |
|  | 8 | -118.4 | \ | \ | \ | \ | \ | \ | 0 |
|  | 9 | -124.1 | \ | \ | \ | \ | \ | \ | 0 |
| Old orchards | 3 | -52.5 | \ | \ | \ | \ | \ | 0–10 | 10 |
|  | 4 | -24.6 | \ | \ | \ | 0–10 | \ | 10–20 | 20 |
|  | 5 | -40.0 | \ | \ | \ | \ | \ | 0–10 | 10 |
|  | 6 | 6.6 | Mild | \ | \ | \ | 0–10, 20–80 | 10–20 | 80 |
|  | 7 | -14.8 |  | \ | \ | 40–60 | 60–80 | 20–40 | 60 |
|  | 8 | -88.3 | \ | \ | \ | \ | \ | \ | 0 |
|  | 9 | -123.8 | \ | \ | \ | \ | \ | \ | 0 |
| Farmlands | 3 | -0.4 | \ |  |  |  |  | 0–10, 40–60 | 30 |
|  | 4 | 0.1 | Mild | 0–10 |  | 10–20 | 20–60 |  | 60 |
|  | 5 | 0.1 | Mild |  |  | 20–60 | 60–80 | 0–20 | 80 |
|  | 6 | 0.8 | Intense | 10–60 | 0–10, 60–80 |  | 80–150 |  | 150 |
|  | 7 | 0.4 | Moderate | 40–60 | 60–80 | 20–40 | 80–100 | 100–150 | 130 |
|  | 8 | -0.9 | \ |  |  |  |  |  | 0 |
|  | 9 | -0.9 | \ |  |  |  |  |  | 0 |

Subsequently, we analyzed soil desiccation at a more accurate scale based on the moisture content of each soil depth. In the young orchards, mildly dry soil layers occurred at the 0–10-cm depth in March, 20–80-cm depths in June, and 20–40-cm depth in July. In addition, moderately dry soil layers occurred at the 0–10-cm and the 40–60-cm depths in June, and not in the 40–60-cm depth in July. In addition, severely dry soil layers were observed at the 0–10-cm depth in April.

In the mature orchards, mildly or severely dry layers were not formed in spring. Dry soil thickness decreased in June, and dry soil layer moved downward to the 60–80-cm depth in July, when compared to the case in the young orchards. The thickness of moderately dry soil layers increased by 10 cm in June and they moved upward to the 20–40-cm depth in July when compared to the young orchards. Severely dry soil layers occurred mainly at the 20– 40-cm (June) and 40–60-cm (July) depths in summer.

Soil desiccation intensity increased in the old orchards when compared to the young and mature orchards. Slightly dry soil layers were observed at the 0–40-cm depths for five months (March–July). The thickness of moderately dry soil layers increased by 10 cm in June, and they moved downward to the 60–80-cm depth in July when compared to those of mature orchards. The occurrence of severely dry soil layers was similar to the case in the young orchard in spring and the mature orchards in summer.

In farmlands, mildly dry soil layers occurred at the 0–10-cm and 40–60-cm depths in March, which shifted to the 0–20-cm depths in May and the 100–150-cm depths in July. Moderately dry soil layers occurred across the 20–150-cm depths across the April–July period. Conversely, severely dry soil layers occurred at the 10–60-cm depths in April–May and contracted to the 20–40-cm depths in July. In addition, intensely dry soil layers were distributed at the 0–10-cm and 60–80-cm depths in June–July. Extremely dry soil layers appeared at the 0–10-cm depth in March and then expanded to the 10–60-cm depths in June– July.

## Discussion

In Weibei rain-fed highland, agricultural production has shifted from cultivating crops to orchard farming, which inevitably changes in soil moisture content. The differences in soil moisture content are mainly associated with spatial differences in root water consumption, plant transpiration, and vegetation canopy on the near-surface evaporation [33]. In the present study, we compared water consumption in the 0–150-cm soil profile between rain-fed wheat fields and apple orchards after farmland conversion in Weibei. The results revealed the underlying water conditions in planted orchards and potential factors mitigating soil desiccation in orchards when compared to farmlands.

### Occurrence of Dry Soil Layers in Orchards and Farmlands

A positive soil water consumption value indicates that soil moisture content has decreased; namely, the water is being consumed at a specific depth interval within the period, which represents a deficit or loss of water; a negative value implies that soil moisture content has increased, o that the water accumulating in a specific soil layer within the period, which represents an increase and water surplus [34]. Here, we analyzed soil water consumption at specific depths and periods in the 0–150-cm profile in orchards and farmlands in Weibei in the course of an apple-growing season. Soil water consumption increased significantly from March through mid-June, along with a rise in temperature and robust physiological activity in apple trees. After the arrival of the rainy season in mid-June, the moisture in the soil in the 0– 150-cm profile was replenished. Based on the total water consumption of the soil profile, the highest soil water consumption in orchards occurred in mid-May to mid-June. According to the results, the soil water consumption in apple orchards was not only closely linked to the evaporation under canopies, but was also correlated with bare ground evaporation between apple tree rows.

Cao et al. [32] calculated soil water over-consumption (*SWO*) as the difference between stable soil moisture content (i.e., water stress point) and forest soil moisture content. The principle is to measure the level of soil water stress on forests based on soil moisture content being below soil water stress point. A positive *SWO* value indicates that the trees are underwater deficiency stress; the higher the value, the more severe the stress. According to the results of the present study, the actual soil moisture content in orchards was always higher than the local stable soil moisture content in Weibei. Therefore, there was almost no water stress in orchard soils, which also means that desiccation was generally absent in the apple orchards.

Here, *SDI* was used as a relatively accurate quantitative indicator of dry soil layers. According to the relevant evaluation thresholds proposed by previous studies [35-36], we analyzed spatiotemporal changes in dry soil layers in apple orchards of different ages in Weibei based on soil depth and time period (month). Dry soil layers were present mainly at shallow depths in the young orchards, which could be related to persistent drought and physiological water consumption by apple trees. The shallow dry soil layers in the young orchards could replenish soil moisture relatively easily, and there were no deep dry soil layers that could not recover moisture easily. Similar results were observed in the mature orchards, with the dry soil layers formed at the shallow depths in close correlation with the persistent drought and the physiological cycles of apple trees. However, the dry soil layers in the old orchards expanded in both time and space compared to those in the young and mature orchards.

Further analyses revealed that the total dry soil thickness reached 60 cm in both the mature and old orchards from mid-June to mid-July, and severe desiccation occurred at specific soil depths. The phenomenon is directly related to the robust physiological activities of apple trees and persistent drought in spring. However, dry soil layers appeared at relatively shallow depths for a short period, which could have influenced the growth and development of apple trees slightly without minimal adverse effects. In addition, the 0–150-cm soil profile in farmlands had five levels of soil desiccation intensity (②–⑥), which indicated that dry soil layers could exist in farmlands not only over long periods, but also across broad spatial ranges and intensities.

### Recovery of Dry Soil Layers in Orchards

Due to poor soil water conditions in Weibei, many researchers consider the region only suitable for the production of herbaceous plants, and that arbor trees may not be grown sustainably in the region. In addition, considering the high transpiration losses via the forest canopy, dry layers could be formed relatively easily in the soil [29]. Other researchers have investigated the causes of dry soil layers on the Loess Plateau, and they have proposed that when the original soil water conditions are poor and reach a certain level of soil desiccation, continuous tree and shrub planting results in continuous soil water consumption, which could worsen the regional soil water conditions [32, 37]. The results of the studies suggest that planting fruit trees in Weibei poses risks with regard to water resource availability.

Notably, we did not observe any permanent dry soil layers in the apple orchards in the present study. Although dry soil layers occurred at particular periods or at specific depths, they could be restored following rainfall. Conversely, different intensities of soil desiccation were observed at the 0–60-cm depths in farmlands, which could not recover moisture easily. First, shading by fruit trees prevents soil water evaporation losses [38]. Secondly, considering the terrain characteristics, the Weibei rain-fed highland is relatively flat. Among the factors influencing soil water balance in the region, the effects of surface runoff and underground runoff are negligible [39]. Furthermore, the soil is deep and thick with groundwater levels at 40–80 m; therefore, there are no groundwater recharge limitations in Weibei. In the absence of irrigation, the soil water input in orchards is limited to natural precipitation, while multiple factors (e.g., root absorption by apple trees, plant transpiration, physiological water consumption, and soil surface evaporation between individual trees) influence the soil water loss. Therefore, planting apple trees does not necessarily lead to the creation of unrecoverable dry soil layers in the rain-fed highland area. The fruit industry could develop sustainably the Weibei area with appropriate orchard management practices.

## Conclusions

This study evaluated soil water consumption characteristics in rain-fed apple orchards and wheat fields based on soil water over-consumption, soil water consumption, and *SDI*. According to the results, dry soil layers occurred at the 0–80-cm depths in the old orchards in mid-June, while they shifted to the 20–60-cm depths in farmlands in May–July. Overall, soil water stress was lower in orchards than in farmlands, and the dry soil layers in orchards could recover. Based on a water consumption perspective, the conversion of wheat fields into apple orchards could mitigate soil desiccation in Weibei.

## Acknowledgments

This work was financially supported by the Fund for Less Developed Regions of the National Natural Science Foundation of China (No. 42167039)

## References

1. Yang WZ, Shao MA. Soil Water Research on Loess Plateau, China (in Chinese). Beijing: Science Press. 2000.

2. Huang MB, Gallichand J. Use of the SHAW model to assess soil water recovery after apple trees in the gully region of the Loess Plateau, China. Agricultural Water Management. 2006; 85(1-2): 67–76.

3. Wang YQ, Shao MA, Zhu YJ, Liu ZP. Impacts of land use and plant characteristics on dried soil layers in different climatic regions on the Loess Plateau of China. Agricultural and Forest Meteorology. 2011; 151(4): 437–448.

4. Zhang YJ. “Dry Belt”: The best witness of rural changes in Weibei (in Chinese). Journal of Western Development. 2019; 259(7): 113–119.

5. Xu M. Study on nutrition diagnosis and evaluation of red Fuji apple leaves in Weibei, Shaanxi Province (in Chinese). Yangling, Northwest A&F University. 2009.

6. Wang XK, Li ZB, Xing YY. Effects of mulching and nitrogen on soil temperature, water content, nitrate-N content and maize yield in the Loess Plateau of China. Agricultural Water Management. 2015; 161(1): 53–64.

7. Wang N, Joost W, Zhang FS. Towards sustainable intensification of apple production in China—Yield gaps and nutrient use efficiency in apple farming systems. Journal of Integrative Agriculture. 2016; 15(4): 716-725 (2016).

8. Zhao ZP, Tong YA, Liu F, Wang XY, Zeng YJ. Assessment of current conditions of household fertilization of apples in Weibei Plateau (in Chinese). Chinese Journal of Eco-Agriculture. 2012; 20(8): 1003–1009.

9. Chen LD, Wei W, Fu BJ, Lü YH. Soil and water conservation on the Loess Plateau in China: review and perspective. Progress in Physical Geography. 2007; 31(4): 389–403.

10. Lü HS, Zhu YH, Skaggs Todd H, Yu ZB. Comparison of measured and simulated water storage in dryland terraces of the Loess Plateau, China. Agricultural Water Management. 2009; 96(2): 299–306.

11. Lü YH, Fu BJ, Feng XM, Zeng Y, Liu Y, Chang RY, Sun G, Wu BF. A policy-driven large scale ecological restoration: Quantifying ecosystem services changes in the Loess Plateau of China. Plos One. 2012; 7(2): 1–10.

12. Cheng LP, Liu WZ, Li Z. Soil water in deep layers under different land use patterns on the Loess Tableland (in Chinese). Acta Ecologica Sinica. 2014; 34(8): 1975–1983.

13. Hu W, Si BC. Revealing the relative influence of soil and topographic properties on soil water content distribution at the watershed scale in two sites. Journal of Hydrology. 2014; 516: 107-118.

14. Ковалев АВ, Билева ОК, Морякова ВК. Комбинированный метод сбора и учета морского зоопланктона... Soviet Journal of Quantum Electronics. 1981; 11: 2486–2492.

15. Li RR, Ma FY, Xiao HL, Wang XP, Kim KC. Long-term effects of revegetation on soil water content of sand dunes in arid region of Northern China. Journal of Arid Environments. 2004; 57(1): 1–16.

16. Mahmood R, Hubbard Kenneth G. Assessing bias in evapotranspiration and soil moisture estimates due to the use of modeled solar radiation and dew point temperature data. Agricultural and Forest Meterology. 2005; 130(1-2): 71–84.

17. Wang YQ, Shao MA, Liu ZP. Large-scale spatial variability of dried soil layers and related factors across the entire Loess Plateau of China. Geoderma. 2010; 159(1-2): 99–108.

18. Yang L, Wei W, Chen LD, Mo B. Response of deep soil moisture to land use and afforestationin the semi-arid Loess Plateau, China. Journal of Hydrology. 2012; 475: 111–122.

19. Zhang YK. Schilling Keith, E. Effects of land cover on water table, soil moisture, evapostranspiration, and groundwater recharge: A Field observation and analysis. Journal of Hydrology. 2006; 319(1-4): 328–338.

20. Giraldo Mario A, Bosch D, Madden M, Usery L, Kvien C. Landscape complexity and soil moisture variation in South Georgia, USA, for remote sensing applications. Journal of Hydrology. 2008; 357(3-4): 405–420.

21. Wang ZQ, Liu BY, Zhang Y. Soil moisture of different vegetation types on the Loess Plateau. Journal of Geographical Sciences. 2009; 19(6): 707–718.

22. Jacobs JM, Mohanty BP, Hsu EC, Miller DA. SMEX02: field scale variability, time stability and similarity of soil moisture. Remote Sensing of Environment. 2004; 92(4): 436–446.

23. Guber AK, Gish TJ, Pachepsky YA, Van Genuchten MT, Daughtry CST, Nicholson TJ, Cady RE. Temporal stability in soil water content patterns across agricultural fields. Catena. 2008; 73(1): 125–133.

24. Brocca L, Melone F, Moramarco T, Morbidelli R. Spatial-temporal variability of soil moisture and its estimation across scales. Water Resources Research. 2010; 46(2): W02516.

25. Biswas A, Si BC. Identifying scale specific controls of soil water storage in a hummocky landscape using wavelet coherency. Geoderma. 2011; 165(1): 50–59.

26. Wang YQ, Shao MA, Liu ZP. Spatial variability of soil moisture at a regional scale in the Loess Plateau (in Chinese). Advances in Water Science. 2012; 23 (3): 310–316.

27. Daniel TK. Microclimate effects of wind farms on local crop yields. Journal of Environmental Economics and Management. 2019; 96: 159–173.

28. Du K, Zhang BY, Li LJ. Soil Water Dynamics Under Different Land Uses in Loess Hilly Region in China by Stable Isotopic Tracing. Water. 2021; 13(2): 242.

29. Yang WZ, Tian JL. Essential exploration of soil aridization in Loess Plateau (in Chinese). Acta Pedologica Sinica. 2004; 41(1): 1–6.

30. Tian L, Zhang JX, Gao JE, Dong JG, Wang YK. Experiment on dry soil water restoration in deep layer (in Chinese). Transactions of the Chinese Society for Agricultural Machinery. 2019; 50(4): 255–262.

31. Bai X, Zhang LH, Wang YB, Tian J, He CS, Liu GH. Varitations of soil moisture under different land use and land cover types in the Qilian Mountain, China (in Chinese). Research of Soil and Water Conservation. 2017; 24(2): 17–25.

32. Cao Y, Li J, Zhang SH, Wang YL, Cheng K, Wang XC, Wang YL, M. Naveed T. Characteristics of deep soil desiccation of apple orchards in different weather and landform zones on the Loess Plateau in China (in Chinese). Transactions of the Chinese Society of Agricultural Engineering. 2012; 28(15): 72–79.

33. Liu BX, Shao MA. Modeling soil–water dynamics and soil–water carrying capacity for vegetation on the Loess Plateau, China. Agricultural Water Management. 2015; 159: 176–184.

34. Qi DL, Hu TT, Song X. Effect of irrigation regime on water consumption pattern and grain yield of seed maize under partial root zone irrigation (in Chinese). Transactions of the Chinese Society of Agricultural Engineering. 2019; (14): 64–70.

35. Yang L, Zhang HD, Chen LD. Identification on threshold and efficiency of rainfall replenishment to soil water in semi-arid loess hilly areas. Science China-Earth Sciences. 2018; 61(3): 292–301.

36. Gou QP, Zhu QK, Li YX, Shen MS, Liu YY, Mei XM, Wang Y. Soil desiccation effects under different vegetation types in the Loess Region of Northern Shaanxi (in Chinese). Acta Ecologica Sinica. 2019; 39(19): 1–8.

37. Li J. et al. Comparison of soil desiccation effects of forest land, grassland and farmland in different precipitation types in the Loess Plateau (in Chinese). Acta Pedologica Sinica. 2008; 45(1): 40–49.

38. Jutamanee K, Onnom S. Improving photosynthetic performance and some fruit quality traits in mango trees by shading. Photosynthetica. 2016; 54(4): 542–550.

39. Jiang DS. Soil Erosion and Control Models in the Loess Plateau (in Chinese). Beijing: China Water Power Press. 1999.

